# Plant Cell Wall Enzymatic Deconstruction: Bridging the Gap Between Micro and Nano Scales

**DOI:** 10.1101/2024.01.11.575220

**Authors:** Yassin Refahi, Aya Zoghlami, Thibaut Viné, Christine Terryn, Gabriel Paës

## Abstract

Understanding and overcoming the resistance of plant cell wall to enzymatic deconstruction is crucial to achieve a sustainable and economical conversion of plant biomass to bio-based products as alternatives to petroleum-based products. Despite the significant scientific advances over the past decades, the plant cell wall deconstruction at cell and tissue scales has remained under-investigated. In this study, to quantitatively characterize plant cell wall deconstruction, we set up an original imaging pipeline by combining time-lapse 4D (space + time) fluorescence confocal imaging, and a novel computational tool, to track and quantify cell wall deconstruction at cell and tissue scales offering a digital representation of cell wall deconstruction. Using this pipeline on poplar wood sections, we computed dynamics of several cellular parameters (e.g. cell wall volume, surface area, and number of cell neighbors) while measuring cellulose conversion. The results showed that the effect of enzymatic deconstruction at the cell scale is predominantly noticeable in terms of cell wall volume reduction rather than a significant decrease in surface area and accessible surface area. The results also revealed a negative correlation between pre-hydrolysis 3D cell wall compactness measures and volumetric cell wall deconstruction. The strength of this correlation was modulated by enzymatic activity. Combining cell wall compactness with the number of neighboring cells as a tissue-scale parameter yielded a stronger correlation. Our results also revealed a strong positive correlation between average volumetric cell wall deconstruction and cellulose conversion, thus establishing a link between key parameters and bridging the gap between nano and micro scales.

## 1 Introduction

The convergence of environmental and economic imperatives due to climate change, increasing global energy demands and unstable oil prices highlights the critical need for a transition from fossil resources towards alternative energy and material sources. Transformation of plant cell wall into bioproducts offers a renewable eco-friendly carbon-neutral alternative to conventional petroleum derived products to achieve sustainable development and to ensure the well-being of future generations [26, 27, 41, 4]. Therefore, transformation of plant cell wall can be regarded as a critical component of a sustainable bioeconomy to meet humanity’s needs while limiting negative impacts on the environment [3]. The primary building blocks of the plant cell wall are cellulose, hemicelluloses, and lignin, which are collectively referred to as lignocellulose. Cellulose is composed of linear chains of D-glucose units linked by *β*-(1,4) glycosidic bonds and is the most abundant biopolymer on Earth [43]. Cellulose fibers are enveloped with interweaving hemicelluloses which constitute a group of heterogeneous polysaccharides made of pentose and hexose sugars and other small groups. Lignin, an irregular phenylpropanoid polymer, forms a protective barrier around the polysaccharides [36, 43, 35]. Although different biotechnological processes are used for lignocellulose conversion, a general process consists of pretreatment of lignocellulosic biomass to improve accessibility of cellulose and hemicellulose [28, 15] for subsequent enzymatic hydrolysis, called saccharification, to break down the carbohydrates into simpler sugars (predominantly glucose) followed by the fermentation of monosaccharides generated by the saccharification [6, 23]. Therefore, enzymatic hydrolysis is a crucial step in the conversion process. A major challenge to enhance the release of fermentable sugars during saccharification is to overcome the natural resistance of plant cell wall to deconstruction, called recalcitrance [18]. The recalcitrance is a major contributor to the high cost of lignocellulosic biomass derived products. Extensive research has been conducted over the past decades to better understand the recalcitrance and its underlying parameters [53, 51, 52]. Several recalcitrance markers like lignin content [39], cellulose crystallinity [16], degree of cellulose polymerization [29], and porosity of the cell wall [17] have been identified. Strikingly, only markers at nano scale have been investigated and plant cell wall enzymatic hydrolysis at cell and tissue scales remains largely under-investigated. Therefore, a notable gap in the literature exists regarding micro scale structural characteristics contributing to recalcitrance. This lack of understanding is mainly due to the lack of comprehensive quantification of cell and tissue scale structural parameters during enzymatic deconstruction. Acquisition and analysis of time-lapse images during enzymatic hydrolysis is particularly challenging, contributing to the deficiency in understanding.

Several methods have been developed to characterize cellular growth and division of plant and animal tissues [13, 48, 32, 14, 37, 49, 22] which can be seen as the opposite mechanism compared to cell wall deconstruction. These innovative and inspiring methods combine confocal time-lapse live imaging and segmentation and tracking. Nevertheless, these methods are not adapted to address the challenges of plant cell wall deconstruction investigation at cell and tissue scales. To acquire time-lapse images of cell wall deconstruction, the cell walls together with an enzyme cocktail should be imaged at relatively high constant optimal temperature (typically 50°C) for enzymatic reaction while keeping the sample stable and avoiding enzymatic cocktail evaporation during hydrolysis for a substantial number of hours. The deconstruction of the cell wall during hydrolysis compromises the suitability of the pre-existing segmentation and tracking methods. In the early stages of cell wall deconstruction, classical methods require time-consuming laborious manual adjustment of parameters, particularly for large datasets. This can introduce bias, compromising the accuracy, reliability, and objectivity of the results. As hydrolysis advances and cell walls develop holes and cracks, possibly culminating in their complete breakdown, these classical algorithms tend to produce increased segmentation errors.

In this study, we develop an original computational pipeline to quantify the evolution of cell and tissue scale structural parameters during plant cell wall deconstruction. Using this pipeline, we quantify dynamics of cell wall structural parameters during enzymatic hydrolysis of poplar wood samples. We also measured the cellulose conversion during the hydrolysis of the poplar samples. Our data revealed that plant cell wall compactness prior to enzymatic deconstruction is negatively correlated with volumetric cell wall deconstruction. Our data also revealed a strong positive correlation between average volumetric cell wall deconstruction and cellulose conversion establishing a link between key parameters and bridging the gap between nano and micro scales.

## Results

### Automated segmentation and tracking of cell wall deconstruction

We selected poplar as our model plant due to its fast growth rate, ease of in vitro cultivation and vegetative propagation, and extensive distribution. Poplar was the first woody plant, the third angiosperm, after *Arabidopsis* and rice, to have its relatively small genome sequenced [29, 33]. We also chose dilute acid pretreatment, one of the commonly used industrial pretreatment processes, because of its relatively low cost and efficiency, particularly with hardwoods [11]. Dilute acid pretreatment solubilizes hemicelluloses, disrupts the lignin structure, and increases cellulose accessibility to enzymes, favoring enzymatic hydrolysis [9, 42]. The chemical composition of the pretreated sample is important for the yield and rate of enzymatic hydrolysis. Therefore, we analyzed the chemical composition of both untreated and pre-treated samples, which indicated a reduction in the fraction of hemicelluloses by approximately 3.2-fold due to the pretreatment (Supplementary Information Appendix, Fig.S1.A1 – 2), in line with previous results [54]. Using plant cell wall natural autofluorescence [10], we collected time-lapse confocal images of poplar wood samples pretreated with dilute acid during enzymatic hydrolysis, following the protocol developed in [55]. Confocal images were acquired every thirty minutes for the first 4 hours, followed by hourly acquisitions for the next 20 hours (Fig.1.A). After acquiring these time-lapse confocal images, we developed an original automated high-throughput 4D image segmentation and tracking pipeline, named WallTrack, to quantitatively characterize cell wall deconstruction. WallTrack simultaneously accomplishes two tasks: i. It computes a 4D voxel (voxel indicates a 3D pixel) resolution map of cell wall deconstruction, despite the movement of the sample during imaging (Fig.1.B), and allows for precise quantification of autofluorescence intensity before and after hydrolysis (Fig. 1.C). ii. It segments individual cell walls and tracks their evolution during enzymatic hydrolysis. WallTrack is devised to address the challenge of cell wall autofluorescence intensity loss during enzymatic hydrolysis, which renders classical segmentation methods unsuitable. Due to the deconstruction of the cell wall, holes and cracks appear in the early hours of enzymatic hydrolysis, potentially leading to the complete disintegration of cell wall segments between neighboring cells. This results in increased segmentation errors such as the fusion of neighboring cells (under-segmentation) or the division of an individual cell wall into multiple smaller, unnecessary segments (over-segmentation) when using classical segmentation methods [13, 48]. Furthermore, enzymatic deconstruction causes separation between the compound middle lamella and the secondary cell wall [54], a process driven by the solubilization and breakdown of cell wall components that act as adhesives between layers [25]. This separation can also lead to over-segmentation errors, where a cell is erroneously divided into two parts. Applying classical methods to datasets with low deconstruction (in early acquisitions with low enzymatic activity) where only cell wall thickness is reduced, requires manual adjustment of the denoising and segmentation algorithms’ parameters to segment each acquisition in the time-lapse datasets. This manual tuning is time-consuming and impractical for large datasets, as each individual acquisition must be checked for segmentation errors. Alterations in parameter values can impact quantifications and therefore can introduce bias, lead to compromised accuracy and reliability, and thereby reduce objectivity. WallTrack addresses these significant limitations by propagating the 3D segmentation of the image before hydrolysis to compute the segmentation of images acquired during hydrolysis. This strategy of propagating spatial information over time, first used on living organisms [2], allows the identification of individual deconstructed cell walls using their earlier state, where the cell walls are intact and the cell boundaries between neighboring cells are clearly marked (i.e. before enzymatic deconstruction). WallTrack first computes the cell wall resolution segmentation of the acquired confocal image before hydrolysis (Fig.2.A). Then, WallTrack registers the acquired confocal image before hydrolysis onto the subsequent confocal images. The computed transformations are then applied to the cell segmentation before hydrolysis to compute segmentations of confocal images acquired during hydrolysis (Fig.2.B, and Supporting Information Appendix Fig.S2). WallTrack automatically generates 3D segmentation of individual cell walls at each time point with unique cell wall identifiers, which remain consistent over time, thereby enabling tracking and analysis of the cell walls during deconstruction.

**Fig. 1:**
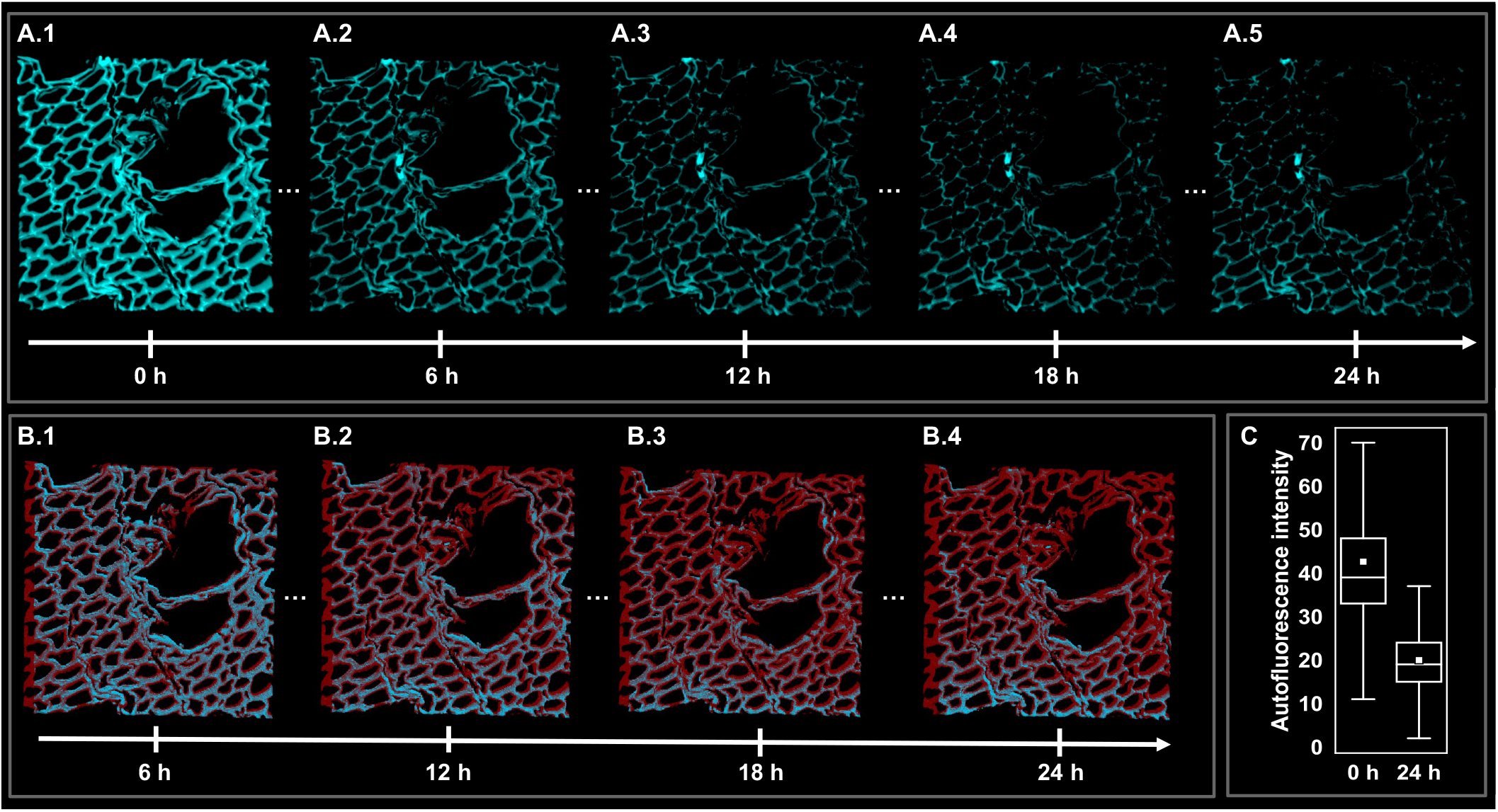
4D imaging of cell wall deconstruction during enzymatic hydrolysis using lignin autofluorescence. (A) Confocal time-lapse images of poplar wood sample were acquired every 30 minutes for the first 4 h followed by acquisitions every hour during enzymatic hydrolysis for the next 20 h (every 6 h is shown) with enzymatic activity of 20 FPU. (B) 4D voxel (voxel is a 3D pixel) resolution map of cell wall deconstruction (color code: red: deconstructed voxels, cyan: remaining voxels) (C) Distributions of cell wall autofluorescence intensity before and after 24 h of enzymatic hydrolysis which show a significant reduction in autofluorescence intensity values (Anova, p-value *<* 0.005). The central lines of the box plot are the median values, and the white squares represent the average values.

### Cell scale impact of enzymatic deconstruction predominantly manifests as a reduction in cell wall volume rather than a reduction in cell wall accessible surface area

To investigate the spatio-temporal cell and tissue scale structural changes during enzymatic deconstruction, an adjacency graph representing the tissular organization and structure was first computed from the segmented 4D images generated by WallTrack. In the adjacency graph, vertices represent the cells and and edges represent cell walls or shared boundaries between neighboring cells (Fig. 3.A). The degrees of the vertices in the adjacency graph were then computed, indicating the number of neighboring cells for each cell, or the number of distinct cell walls that are shared with adjacent cells [24, 5]. For an individual cell, the number of cell neighbors is equivalent to the number of cell junctions which exhibit a unique composition compared to other parts of the cell wall, typically having a higher lignin content [51, 1]. After determining the adjacency graph, we focused on the cell wall volume and cell wall surface area as two key indicators of size at cell scale. This focus was prompted by visual observations of individual cells during hydrolysis, which highlighted changes in cell volumes and cell wall surface areas (Fig. 2.B.1 – B.5 and Fig. 3.C & D). Thus, we computed the individual cell wall volumes using the segmented time-lapse images. We observed a significant reduction of the cell wall volume before and after 24 h of hydrolysis for all time-lapse datasets collected in the presence of enzymes (Anova test, p-value *<* 0.005) (Fig. 3.B.1 & E). We then computed the total surface area of the cell walls (Fig. 3.B.2 & E). The surface area directly influences the enzyme-substrate interactions since enzymes require adequate access to their target substrates within the complex lignin-embedded polysaccharide matrices [47]. We observed a general reduction in the total surface area, however, we did not find a conclusive statistically significant difference between the surface area before and after 24 h of enzymatic deconstruction. We then computed the accessible surface area (ASA) by taking into account the wall surface regions which are directly in contact with enzymes and discarding the cell walls shared by neighboring cells. The cell wall accessible surface area can be seen as a cell scale counterpart to the accessible surface area of the cell wall polysaccharide matrix (the higher accessible cell wall surface, the higher accessibility to polysaccharides) and is one of the most important parameters affecting the plant cell wall enzymatic deconstruction yield and rate [30, 31]. Similar to the total surface area, a reduction was observed in the accessible surface area which was not statistically conclusive (Fig. 3.B.3 & E). We consequently chose the cell wall volume, the 3D space occupied by the lignocellulosic constituents of the cell, and its evolution during enzymatic deconstruction as the size metric providing a quantifiable cell scale indicator to characterize the enzymatic deconstruction of the cell wall.

**Fig. 2:**
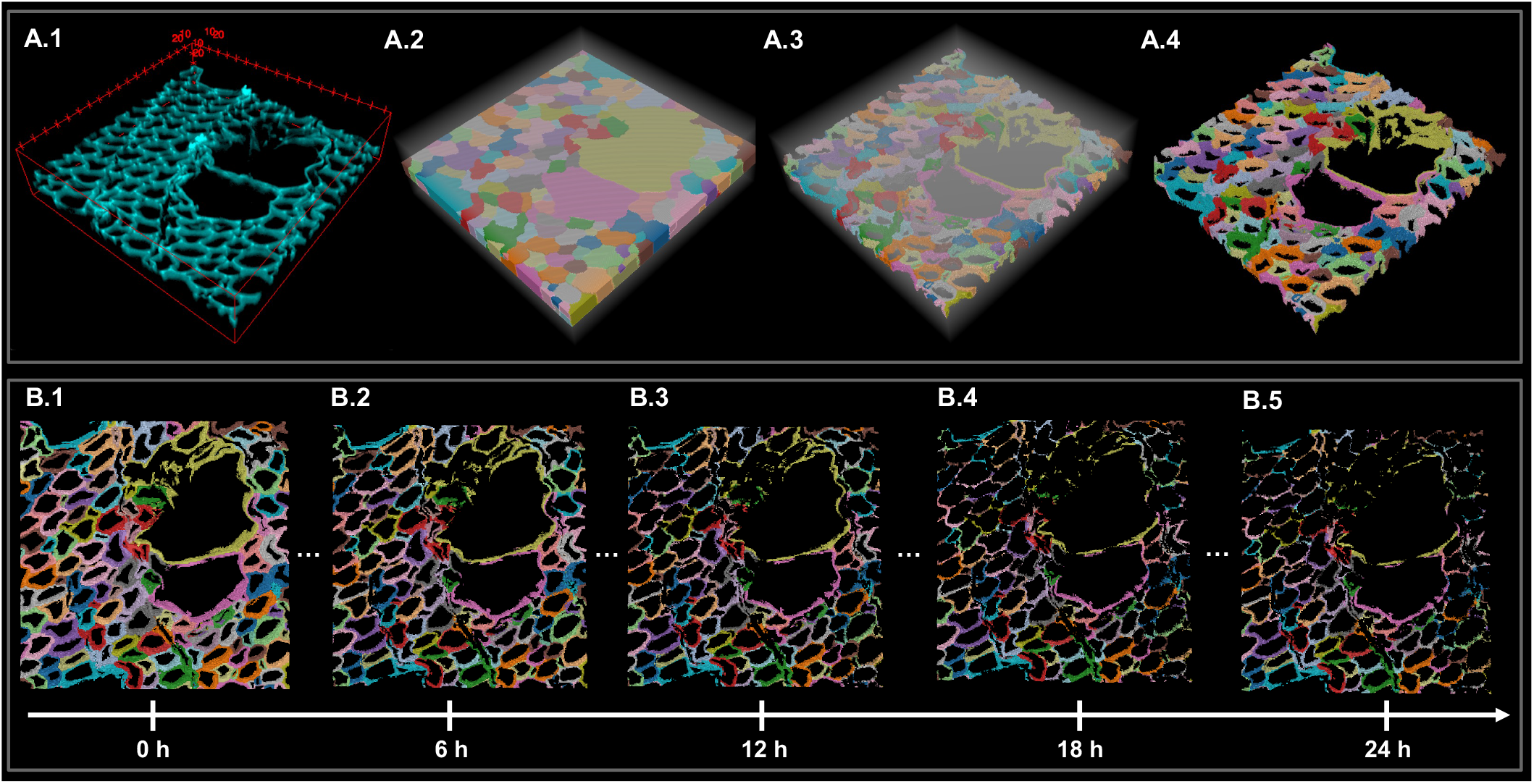
4D segmentation and tracking of cell walls during enzymatic hydrolysis. (A) Segmentation of the image before hydrolysis. (A.1) Confocal acquisition before enzymatic hydrolysis (A.2) Cell resolution segmentation of the image before enzymatic hydrolysis obtained using watershed algorithm with auto seeding. The background is marked by a semi-transparent white color. (A.3) Cell wall segmentation obtained using thresholding. (A.4) Background removal. The cells and cell walls are colored randomly to facilitate the visual distinction of neighboring cells. (B) Cell wall segmentation and tracking of the confocal time-lapse images shown in (Fig. 1.A) obtained using the WallTrack pipeline (every 6 h is shown). Cell walls are colored according to lineages.

**Fig. 3:**
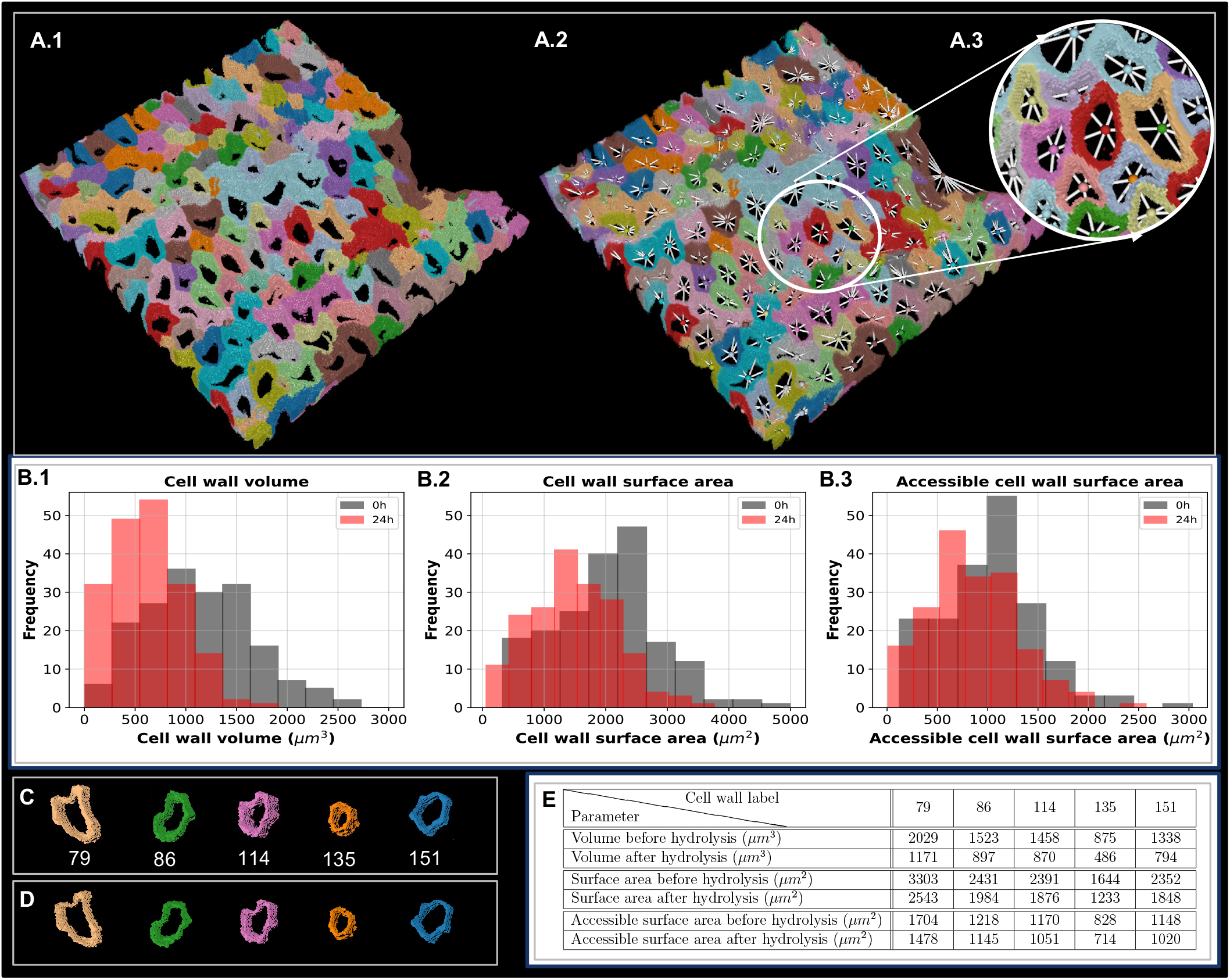
Quantification of cell and tissue scale parameters during enzymatic hydrolysis. (A) Extraction of adjacency graph from segmented images. Cell adjacency graph represents the tissue’s structure and its intercellular connections by a graph computed from the segmented image (A.1). Cells are represented as nodes (vertices), and shared cells wall are represented by edges between neighboring cells (A.2) in the adjacency graph. (B) Cell and tissue scale parameter quantification before and after hydrolysis (B.1) Cell wall volume distribution before and after 24 h of hydrolysis. We can observe a statistically significant shift to the left of the distribution of cell wall volumes after hydrolysis (Anova test, p-value *<* 0.005, N = 184) (B.2) Distributions of cell wall surface areas of individual cells before and after 24 h of hydrolysis (N = 184). (B.3) Distributions of accessible cell wall surface areas before and after 24 h of hydrolysis (N = 184). (C) Randomly selected cell walls before hydrolysis with their unique identifiers (labels) shown in the row directly below. (D) The selected cell walls during hydrolysis which exhibit changes due to enzymatic hydrolysis (E) Cell wall volume, surface area, and accessible surface area before and after hydrolysis for the selected cells shown in C and D.

### Three dimensional cell wall compactness measures prior to hydrolysis are correlated with volumetric cell wall deconstruction whose strength is modulated by the enzymatic activity

Building upon the quantification of cell wall volume and cell wall surface area, we further investigated the cell and tissue scales parameters to study whether they can be informative about the enzymatic deconstruction at cell and tissue scales. More precisely, we studied whether the pre-hydrolysis cell and tissue scale parameters (i.e. parameters measured prior to enzymatic deconstruction) can potentially serve as predictive markers for volumetric cell wall deconstruction. The volumetric cell wall deconstruction after 24 h of enzymatic hydrolysis, denoted hereafter by *D*^24*h*^, is defined as 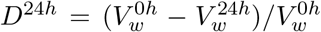, where 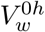 and 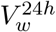 are cell wall volumes before and after 24 h of enzymatic deconstruction respectively. We first computed the Pearson correlation between cell wall volume before hydrolysis, 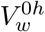, and *D*^24*h*^, for the datasets collected in the absence (0 FPU) and in the presence of enzymes with three different enzyme loadings of 5, 10, and 20 FPU/g of glucan (Fig. 4.A.1 – 4). This range of enzymatic activities was selected to represent varying levels and to examine their effects on cell wall deconstruction. We found a very weak positive correlation for datasets collected with the highest enzymatic activity value (20 FPU), and a very weak negative correlation for the datasets collected in the absence of enzymes, with no significant correlations for datasets collected with other enzymatic activities (5 and 10 FPU). We then computed the Pearson correlation between the number of cell neighbors and *D*^24*h*^. A weak negative correlation was observed exclusively for the datasets collected with enzymatic activity value of 20 FPU. To investigate the relationship between surface area and volumetric cell wall deconstruction, the Pearson correlations were then computed between *D*^24*h*^ and the total cell wall surface area and cell wall accessible surface area before deconstruction. We found weak positive correlations for the highest enzymatic activity values (10 and 20 FPU) for both pre-hydrolysis total cell wall surface area and accessible cell wall surface area, with higher correlation coefficients for pre-hydrolysis accessible surface area compared to the coefficients computed for pre-hydrolysis total surface area. To sum up, the correlations between initial cell wall volume, number of adjacent cells, cell wall surface area, and accessible cell wall surface area were not conclusively correlated with volumetric cell wall deconstruction. Consequently, we redirected our focus to investigate the relationship between the compactness of the cell wall before hydrolysis, quantified as a ratio of cell wall volume to cell wall surface area, and its volumetric deconstruction. The cell wall compactness at level of polymers’ matrix is essentially connected to the density and arrangement of polymers (mainly cellulose, hemicelluloses, and lignin) within the cell wall structure. Consequently, compactness is linked to the restricted enzymatic accessibility to polysaccharide fibers due to the tight packing of the wall components [29, 53]. We therefore first computed the cell wall Compactness Index (*CI*), a dimensionless parameter which is the ratio of volume over the total surface area raised to the power of three-halves (Fig. 4.C). The definition of *CI* implies that if two cell walls have the same volume, the one with a smaller surface area is more compact. We found significant negative correlations between *CI* and *D*^24*h*^ in datasets collected in the presence and absence of enzymes (Fig. 4.A.1 – 4). We then computed the Accessible Compactness Index (*ACI*) where the accessible cell wall surface area is used rather than the total surface area to determine the cell wall compactness (Fig. 4.C). We found significant negative correlations between *ACI* and *D*^24*h*^ in the presence and absence of enzymes with higher correlation coefficients compared to those obtained between *CI* and *D*^24*h*^. Building on this result, we further refined our approach by incorporating the number of cell neighbors into the compactness calculation, a parameter that we called Compactness Neighbor Index and denoted by *CNI*. This integration aimed to account for the potential influence of surrounding cells on the deconstruction process, hypothesizing that the local cellular environment might play a role in determining the susceptibility of the cell wall to enzymatic deconstruction. The rationale behind this was that the number of neighboring cells could be indicative of the compactness at the tissue scale, providing insights into how closely packed cells within a tissue can affect enzymatic access and efficiency. *CNI* was computed by multiplying *ACI* by the square root of the number of neighboring cells. Remarkably, the inclusion of the number of cell neighbors into our compactness metric yielded even more pronounced results. We observed a higher correlation between *CNI* and volumetric cell wall deconstruction compared to *ACI* alone (Fig. 4.A.1 – 4). This enhanced correlation indicates that *CNI* appears to be a more reliable predictor of volumetric cell wall deconstruction and underlines the importance of the intercellular context in which enzymatic deconstruction occurs. Furthermore, the comparison of correlation coefficients of different parameters computed for the data collected in the absence of enzymes and in the presence of enzymes with different activities revealed that the strength of correlations is modulated by the level of enzymatic activity such that the strength of the correlation coefficients increased with higher enzymatic activity. The difference between the correlation coefficients between the control datasets and the datasets collected with enzymatic activity of 5 FPU was not significant. We then computed the Pearson correlation between cell wall volume, cell wall surface area, number of neighbors, *ASA, CI, ACI, CNI* prior to deconstruction and volumetric cell wall deconstruction during hydrolysis, i.e. 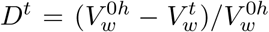, 0*h < t* ≤ 24*h* where 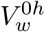 and 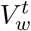 are cell wall volumes before and after *t* h of enzymatic deconstruction respectively (Fig. 4.B.1 – 4). The results revealed the same statistical trends with no conclusive correlations between *D*^*t*^ and the pre-hydrolysis cell wall volume, number of cell neighbors, cell wall surface area, and accessible cell wall surface area. We also observed that the negative significant correlations between pre-hydrolysis compactness parameters (i.e. *CI, ACI*, and *CNI*) and volumetric cell wall deconstruction were consistent across different time intervals within the 1 to 24 h range of enzymatic deconstruction. The higher enzyme activities consistently led to stronger correlations for *D*^*t*^, 1 *< t* ≤ 24*h*. This consistency across various time points underscores the robustness of the cell scale compactness metrics, *CI, ACI*, and *CNI*, in capturing key structural determinants of cell wall deconstruction over the course of enzymatic activity.

**Fig. 4:**
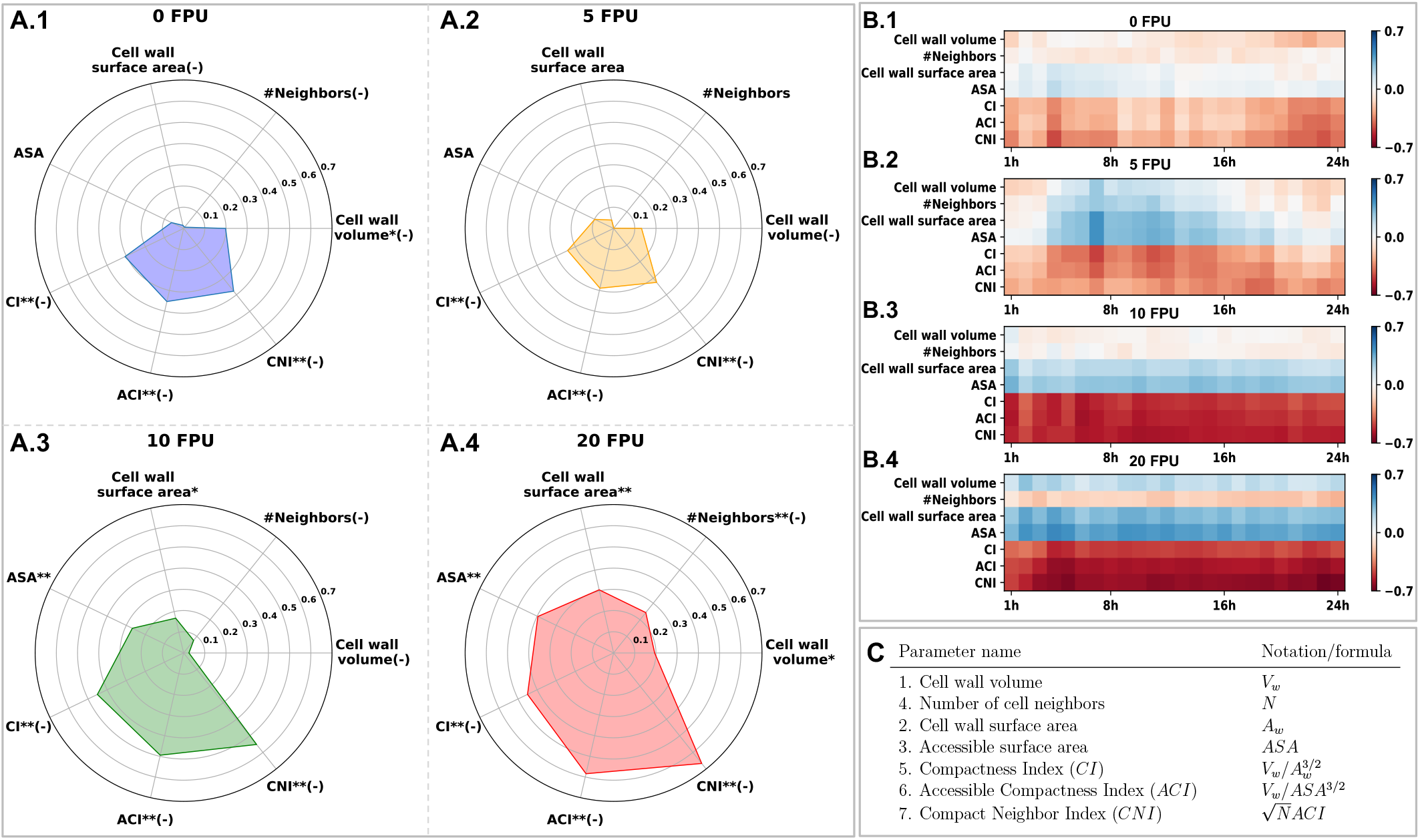
Correlation coefficient values between cell and tissue scale parameters prior to enzymatic deconstruction and volumetric cell wall deconstruction. (A.1) Pearson correlation coefficient values between volumetric cell wall deconstruction after 24h of enzymatic hydrolysis and initial cell and tissue scale parameters in the absence of enzymes. (A.2 – 4) Pearson correlation coefficient values between volumetric cell wall deconstruction after 24 h of enzymatic hydrolysis and pre-hydrolysis cell and tissue scale parameters in the presence of enzymes with enzymatic activity of 5, 10, and 20 FPU respectively. (B.1) Correlation matrix showing correlation coefficients between volumetric cell wall deconstruction during enzymatic hydrolysis for 24 h and initial cell and tissue scale parameters in the absence of enzymes (0 FPU). (B.2 – 4) Correlation matrix showing correlation coefficients between volumetric cell wall deconstruction during enzymatic hydrolysis for 24 h and pre-hydrolysis cell and tissue scale parameters in the presence of enzymes for enzymatic activities of 5, 10, and 20 FPU respectively. (C) Table of notations and formulas for cell and tissue scales parameters. Accessible Surface Area (ASA) within the table and sub-figures denotes the accessible cell wall surface area. (-) denotes negative correlations and statistical significance of correlation coefficients are marked with * and ** for p-value *<* 0.05 and p-value *<* 0.005 respectively. Numbers of individual cells before hydrolysis are 185, 177, 205, and 202 for datasets collected with enzymatic activity of 0FPU, 5FPU, 10FPU, and 20FPU respectively. Total number of cells in the time-lapse datasets analyzed in B.1 – B.4 are 5180, 4956, 5740, and 5656 for datasets collected with enzymatic activity of 0 FPU, 5 FPU, 10 FPU, and 20 FPU respectively.

### Average volumetric cell wall deconstruction is strongly correlated with cellulose conversion

To elucidate the connection and interplay between parameters at the polymer, and cell scales, our study focused on analyzing cellulose conversion in poplar sections within mini-reactors over a 24-hour period and we determined the cellulose conversion of poplar sections during 24 h with three enzyme loadings of 5, 10, and 20 FPU/g of glucan. Our measurements indicated a marked increase in cellulose conversion during the initial 4 hours, followed by a more gradual rise in the subsequent 20 hours, suggesting a slowdown in the enzymatic process (Supporting Information Appendix, Fig.S1.B). We also observed that higher enzyme loading levels corresponded to increased rates of cellulose conversion. In contrast, experiments conducted in the absence of enzymes yielded significantly lower cellulose conversion rates. These results were in line with the expected outcomes and previous research [21, 19]. We then sought to establish a quantitative relationship between the cellulose conversion and the volumetric cell wall deconstruction. For this purpose, we computed the Pearson correlation coefficients between the cellulose conversion and the average volumetric cell wall deconstruction during hydrolysis for different enzymatic activities (0, 5, 10, and 20 FPU). We found a highly significant, strong positive correlation (correlation coefficient ¿ 0.98, p-value *<* 0.005) between cellulose conversion rates and the average volumetric cell wall deconstruction for all enzymatic activities. This significant correlation underscores a quantifiable linear relationship that bridges molecular-level cellulose conversion with cell and tissue scales wall deconstruction. Moreover, this result demonstrates that the volumetric deconstruction of cell walls is a relevant indicator of the deconstruction of the plant cell wall into glucose. This aspect is particularly crucial in the context of the saccharification process, where the efficiency of converting plant biomass into simple sugars is of paramount importance.

## Discussion

In this study, we characterized the plant cell wall deconstruction at cell and tissue scales by devising a novel computational pipeline to segment and track the cell walls, allowing the assessment of morphological parameters’ evolution during enzymatic hydrolysis. Several studies have successfully integrated various imaging techniques with segmentation and tracking algorithms to characterize cell growth and division in plant and animal tissues [48, 37, 14, 12]. However, these methods are not adapted to 4D cell wall deconstruction datasets because of the reduction in the cell wall autofluorescence intensity due to the change in lignin environment occurring during enzymatic hydrolysis. During the initial phase of cell wall enzymatic hydrolysis, where only the cell wall thickness is reduced, classical methods require manual adjustment of denoising and segmentation algorithm parameters to segment each time-lapse acquisition. This manual tuning is laborious, time-consuming and impractical for large datasets, requiring individual checks for segmentation errors. In addition, modifying these parameters may impact the quantitative results, possibly leading to the introduction of bias, reducing accuracy and reliability, and thus compromising objectivity. As the hydrolysis progresses, holes and cracks begin to appear in the cell walls, potentially followed by complete deconstruction of cell walls. This leads to segmentation errors such as the merging of neighboring cells (under-segmentation) or the division of a cell into two parts (over-segmentation), when classical algorithms are used. WallTrack addresses these drawbacks by initially segmenting images before enzymatic deconstruction, at a stage where cell walls are intact and the boundaries between adjacent cells are distinct. This initial segmentation is then propagated over time to compute the segmentation of images acquired during hydrolysis. Thereby, WallTrack provides an automated high throughput 4D imaging pipeline devised to segment and track lignocellulose deconstruction at cell wall resolution leading to a detailed tissue-wide virtual representation of plant cell wall deconstruction. Therefore, WallTrack can be considered as a significant step towards devising a digital twin of lignocellulose deconstruction which will incorporate the 4D virtual tissue provided by WallTrack as the digital representation and accurate mapping of real-world counterpart [46, 50].

The quantification of cell wall volume, surface area, and accessible surface area before and after 24 hours of enzymatic deconstruction revealed that, although there was an overall decrease, the change in surface area was not statistically significant across all datasets. However, a significant decrease in cell wall volume was consistently observed across all datasets after 24 hours of hydrolysis. This indicates that the effect of enzymatic deconstruction at the cell scale is predominantly noticeable in terms of cell wall volume reduction and enzymatic hydrolysis has a more uniform and pronounced impact on the cell wall volume.

We showed that pre-hydrolysis 3D cell wall compactness is negatively correlated with volumetric cell wall deconstruction and the strength of the correlations is modulated by the enzymatic activity (Fig. 4). Among the parameters that we investigated, we found a weak significant correlation between cell wall surface area exclusively for the datasets collected with the highest enzymatic activities (10 and 20 FPU). Similar results were obtained for the accessible cell wall surface area. For the cell wall volume, we did not get conclusive correlations. This indicates that cell wall compactness, as a parameter combining cell wall volume and cell surface area, is a more reliable indicator of volumetric cell wall deconstruction compared to the cell wall volume and cell wall surface area. A more compact cell wall can be more resistant to pretreatment and enzymatic deconstruction because the tight packing of cellulose, hemicelluloses, and lignin makes it more difficult for enzymes to access and break down these polymers [54]. Our results show that utilizing the accessible surface area rather than the total surface area to compute cell wall compactness (*ACI*) increased the strength of correlations with volumetric cell wall deconstruction. The accessible surface area represents the actual area, or portions of the cell wall, available for enzymes to work effectively. Compared to the total cell wall surface area, the accessible surface area provides a more relevant measure of cell wall material which is available for the action of enzymes due to the numerous cell contacts in plant tissues. Our results showed that the number of cell neighbors is negatively correlated with the volumetric cell wall deconstruction exclusively for the highest value of enzymatic activity. A higher number of cell neighbors implies a higher number of lignin-concentrated recalcitrant cell junctions [51, 1] and supports the inverse relationship between the number of neighbors and volumetric cell wall deconstruction. Consequently, the integration of the number of neighbors and cell wall compactness, *CNI* parameter, leads to stronger correlations regardless of level of enzymatic activity. This provides a single dimensionless parameter integrating both cell scale (compactness) and tissue scale parameter (number of neighbors) as an indicator of volumetric cell wall deconstruction. The use of the square root transformation suggests that while the presence of neighboring cells impacts deconstruction, this impact does not increase linearly with their number and the impact of each additional neighbor becomes progressively smaller, underscoring a non-linear relationship. Dimensionless parameters are highly relevant because they allow comparison of results across different time-lapse datasets which can have different sizes and dimensions and facilitate the scaling of computational results to real-world, full-scale applications, ensuring that derived insights are applicable across various sizes and conditions. Moreover, we evidenced a more intricate relationship between cell wall compactness measures and volumetric cell wall deconstruction where the strength of the correlation is modulated by the level of enzymatic activity where a higher level of enzymatic activity corresponds to an increased correlation coefficient. This shows that the level of enzymatic activity clarifies and enhances the relationship between key parameters. Thus, in comprehensive predictive analysis of tissue scale cell wall deconstruction, the consideration of level of enzymatic activity is critical.

Importantly, our approach revealed a strong and compelling correlation between average volumetric cell wall deconstruction and cellulose conversion. This finding indicates a measurable, linear connection that links the molecular-level transformation of cellulose to changes at the cell scale in cell wall volume. Moreover, this outcome underscores that the volumetric deconstruction of cell walls is a relevant indicator of the deconstruction of the plant cell wall into glucose. This aspect is particularly crucial in the context of the saccharification process, where the efficiency of converting plant biomass into simple sugars is of paramount importance. The establishment of this quantitative relationship across scales underscores the benefit, impact and significance of investigating enzymatic deconstruction at cell and tissue scale and paves the way for further studies into plant cell wall deconstruction at these specific scales. This result advantageously completes previous studies aiming at deciphering the mode of action of enzymes, particularly cellulases at the nanoscale, with the limitation that only model cellulose was used [7, 20]. While our study is specific to pretreated poplar wood sections and further research is required to assess the applicability of our findings to different biomass and pretreatment combinations, it represents first evidence of a quantitative link between nano- and macro-scale markers on real biomass samples. Since the enzymatic hydrolysis is a critical step in the conversion of plant cell wall into bio-based products, the finding of this significant robust relationship paves the way to better understand the saccharification. Indeed, synergy and interplay between hydrolases and oxidases such as LPMOs acting on lignocellulose is still under debate considering optimal partnering [44] while spatial localization of LPMO action at the cellular scale has been rarely investigated [8]. Overall, the strategy we have set up should contribute to open and develop new paths towards these fundamental enzymatic questions.

Global challenges like climate change, looming scarcity of fossil resources together with growing world population, require that we innovate to improve the way we produce and consume to limit our negative impact on the environment [45]. Biomass can play an important role in reaching the global climate objectives set in the Paris Agreement on climate change [38]. Our study provides quantitative novel insights into enzymatic deconstruction at under-investigated cell and tissue scales and bridges the gap between polymer and tissue scales. This study is a fundamental step towards comprehensive understanding of enzymatic hydrolysis yield. In the long run, our approach can directly influence the economic and operational feasibility of biorefineries and should contribute to sustainable energy strategies, material development, and mitigating environmental impact.

## 2 Materials and Methods

### Sample preparation and compositional analysis of biomass

Poplar stems (*Poplus nigra x deltoides*) were cut into fragments of 0.4 cm wide, 2 cm long and 0.2 cm thickness using a rasor blade. Poplar fragments were then pretreated with 2% of dilute sulfuric acid at 170°C during 20 minutes following the protocol in [54]. Fragments of native and pretreated poplar were then prepared from transverse sections of xylem as 40 *μ*m thickness using a sliding microtome (Stemi 1000, Zeiss, Germany). To perform compositional analysis, the samples were milled into 80 *μ*m size particles. Compositional analysis including moisture, ash, lignin, and carbohydrates’ contents of native and pretreated poplar samples performed using the method described in [17].

### Enzymatic saccharification

Enzymatic hydrolysis assays on poplar sections was performed with a commercial preparation Cellic CTec2^®^ (Novozymes A/S Bagsværd, Danemark), selected for its hydrolysis efficiency with a cellulase activity of 195 FPU/mL.

For validation purposes, enzymatic hydrolysis were performed in two different systems:

- in a customized incubation chamber containing 60 *μ*L of acetate buffer and 0.02% of sodium azide and the Cellic CTec2 ^®^ cocktail with a final enzyme concentration of 40 FPU/g of glucan for 24 h at 50°C without agitation to perform hydrolysis conditions identical to those between a slide and a cover slip under the microscope. The reaction mixtures (buffer and sodium azide) with the poplar sections were pre-incubated for 30 min at 50°C. The enzymatic hydrolysis was then initiated by adding the enzyme cocktail.
- mini reactors made of embedded capsules of 6 mm diameter with snap-on integral caps made from polypropylene, highly-resistant to high temperature, in which the same volume of the reaction mixture as for the sealed frame was used.

After 24 h of enzymatic hydrolysis, the reaction mixture was analyzed and the concentration of glucose released from enzymatic hydrolysis in the supernatant was determined using a high-performance anionic exchange chromatography (HPAEC-PAD, Dionex) to calculate the cellulose conversion from the pre-treated poplar, as previously described [17]. All experiments and analysis were carried out as triplicates.

### Customized incubation chamber for imaging during enzymatic hydrolysis

40 *μ*m-thick cross sections prepared from both native and pretreated poplar samples were incubated in 0.05 M acetate buffer at pH 5 for 30 min prior to mounting in the same medium on a microscope slide covered with a coverslip (0.17 mm thickness) separated by a spacer (Gene frame ^®^ 65 *μ*L, Thermo Scientific) to provide a 65 *μ*L sealed chamber containing 60 *μ*L enzyme-buffer solution to avoid the evaporation during the enzymatic hydrolysis. The samples were fixed to the cover slide using a temperature resistant adhesive to avoid the movement of the samples in the reaction mixtures. Optimal temperature was maintained at 50°C using a microscope adapted incubator (H301-Mini-OKOLab, Italy).

### Fluorescence confocal time-lapse acquisition protocol

To acquire time-lapse confocal images, we used a laser scanning microscope (Leica TCS SP8, Germany) equipped with 63× oil-immersion objective (NA = 1.4), z-stacks (with 0.3*μ*m z-step) of both native and pretreated poplar samples were acquired at scan speed of 400 Hz with a resolution of 256 × 256 pixels. A 405 nm laser (4% intensity) was used for imaging cell wall sample autofluorescence by detecting fluorescence emission on the 415-700 nm range using the HyD detector in counting mode. Confocal z-stacks of native and pretreated poplar were taken every 30 min for the first four hours and every one hour until 24 h. The time interval, image resolution and microscope parameters were optimized to avoid sample photo-bleaching. The optimal microscope parameter values were determined as a compromise between suitable acquisition quality for subsequent segmentation and reduced laser exposure.

### WallTrack

Let {*I*^0^, …, *I*^*T*^ } be the time-lapse 3D confocal images where *I*^*t*^, is the 3D confocal image (z-stack) acquired at time point *t*, 0 ≤ *t* ≤ *T, T* is total time of data acquisition which is 24 h in this study. The z-stacks included empty slices together with some noise on their top and bottom sections with the imaged poplar sample in the middle section. Special care was taken to have the sample surface normal as close as possible to Z direction. To segment *I*^0^, in the main text referred to as the image before hydrolysis, where the cell walls are intact and not subjected to enzymatic deconstruction, WallTrack first computes the average signal intensity per voxel (volumetric pixel) for each slice of the *I*^0^. To identify the poplar sample in *I*^0^, WallTrack dismisses top and bottom slices whose average signal intensity per voxel values were below or equal to a threshold (a unique threshold value (= 8) defined manually) and applies a 3D watershed algorithm [48] of the middle section after denoising using Gaussian filter and alternative-sequential filter (ASF). WallTrack computes the watershed seeds using the h-minima operator by identifying local minima regions [48]. WallTrack computes the cell wall segmentation of *I*^0^, denoted by *S*^0^, using thresholding, as follows:

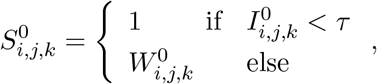

where *W* ^0^ is the watershed segmentation, and *i, j, k* ∈ ℕ, *i* ≤ *M, j* ≤ *N, k* ≤ *K, M, N, K* are X-Y resolution, and number of image slices of *I*^0^ respectively. *τ* is a global threshold value selected to minimize the signal loss in Z direction. 1 is the background label in the segmented images.

To segment the z-stacks during enzymatic hydrolysis, (i.e. *I*^*t*^, 0 *< t* ≤ *T* ), WallTrack first computes the affine transformation that linearly registered *I*^0^ onto *I*^*t*^ using block matching algorithm [34]. WallTrack uses the affine transformation to initialize the block matching algorithm to compute the non-linear transformation 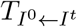 to register *I*^0^ onto *I*^*t*^. Thresholding was then used to calculate the cell wall segmentation of *I*^*t*^denoted by *S*^*t*^ following the application of the computed nonlinear transformation to *W* ^0^, (i.e. 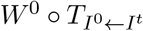):

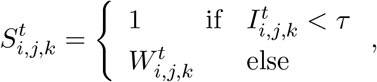

where 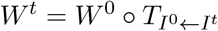, *i, j, k* ∈ ℕ, *i* ≤ *M, j* ≤ *N, k* ≤ *K, M, N, K* are X-Y resolution, and number of image slices of *I*^*t*^ respectively (Supporting Information Appendix Fig.S2). To compute cell wall volume, voxel counting method was used where the number of voxels having the same label were computed and was multiplied by the voxel volume which was approximately 0.155 *μm*^3^ due to the imaging resolution. The cells at the margins of the images together with small cell segments (whose volumes are typically smaller than 400 *μm*^3^) which comprises the precision of quantification because of the movement of the sample during acquisition are discarded from the analysis. To compute cell wall surface area triangular meshes of cell wall surfaces were generated using the marching cubes algorithm from the Visualization Toolkit (VTK) [40]. To compute cell surface area triangular meshes were generated from watershed segmentations. The accessible surface area, *ASA*, was computed as *ASA* = (*A*_*w*_ + *A*_*l*_ − *A*_*c*_)*/*2, where *A*_*w*_, *A*_*l*_, and *A*_*c*_ are cell wall surface area, lumen surface area, and cell surface area respectively. WallTrack is implemented in Python 3 language using Numpy, Scipy, and matplotlib packages.

## Data Availability

WallTrack code and data analysis scripts to generate the figures with associated data are available on demand and will be available upon acceptance via the FARE laboratory GitLab repository (https://gitlab.com/farelab/teamyr/publications/refahi_et_al_4d).

## Acknowledgments

We thank Anouck Habrant for her help in confocal imaging. We thank Grégoire Malandain, Solmaz Hossein Khani, Khadidja Ould Amer, and Ali Faraj for their comments on the manuscript. This work was supported by Agence Nationale de la Recherche (ANR) through “BIOMOD” (ANR-19-CE43-0010) grant to Y.R, and by Grand Est Region and FEDER through “TECMI-4D” PhD funding to A.Z.

## Author contribution

Y.R., A.Z, C.T, and G.P. designed research; Y.R., A.Z., and T.V. performed research; Y.R. and T.V. analyzed data; Y.R., and G.P. wrote the paper. All authors read and approved the manuscript.

