## Supporting Information for "Plant Cell Wall Enzymatic Deconstruction: Bridging the Gap Between Micro and Nano Scales"

**This PDF file includes:**

Figs. S1 to S2

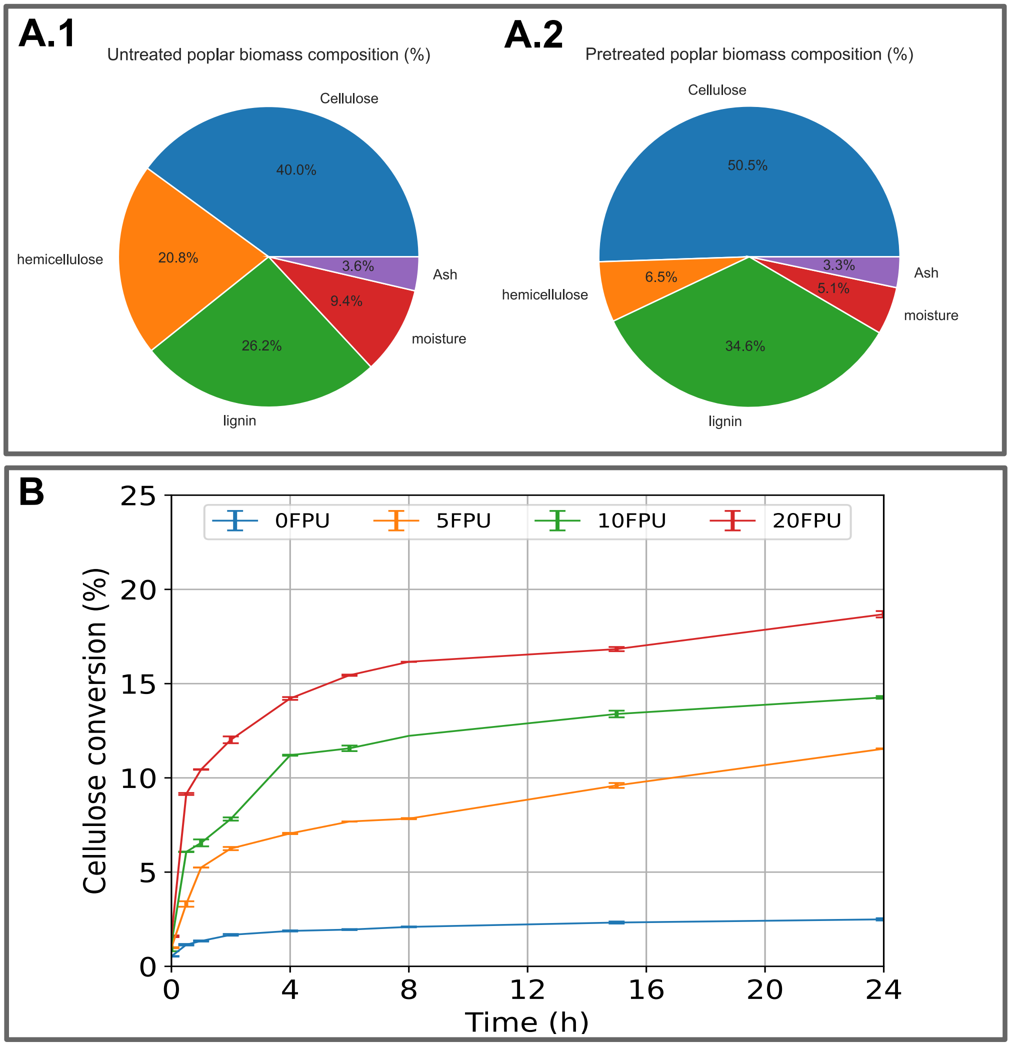


Fig.S 1: (A) Chemical composition of untreated and pretreated poplar samples. (B) Cellulose conversion of four datasets collected with enzymatic activity of 0 FPU (absence of enzymes), 5 FPU, 10 FPU, 20 FPU.

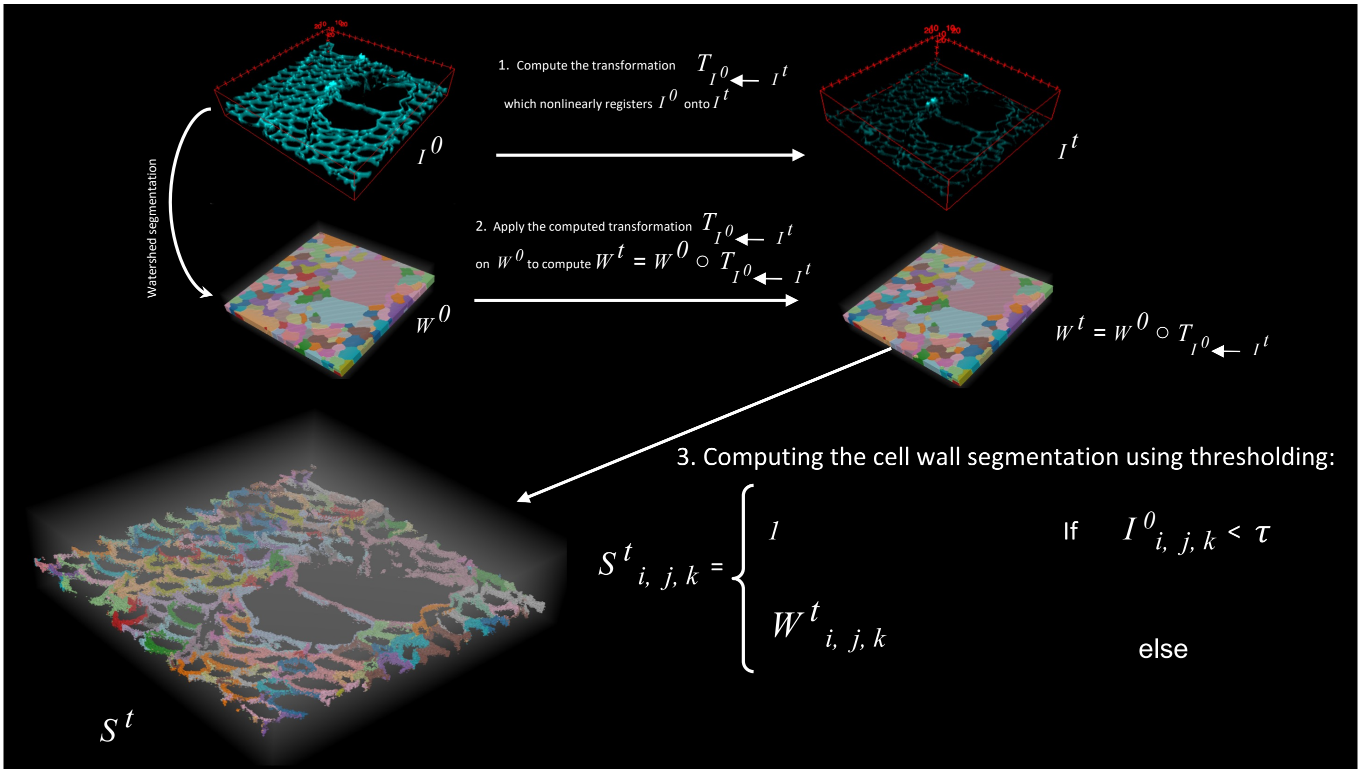


Fig.S 2: WallTrack registers the acquired confocal image before hydrolysis onto the following confocal images. The computed transformations are then applied to the cell segmentation before hydrolysis to compute segmentations of confocal images acquired during hydrolysis. The backgorund is marked by 1.
